# Coherent synthesis of genomic associations with phenotypes and home environments

**DOI:** 10.1101/051110

**Authors:** Jesse R. Lasky, Brenna R. Forester, Matthew Reimherr

## Abstract

Local adaptation is often studied via 1) multiple common garden experiments comparing performance of genotypes in different environments and 2) sequencing genotypes from multiple locations and characterizing geographic patterns in allele frequency. Both approaches aim to characterize the same pattern (local adaptation), yet the complementary information from each has not yet been coherently integrated. Here, we develop a genome-wide association model of genotype interactions with continuous environmental gradients (G×E), *i.e.* reaction norms. We present an approach to impute relative fitness, allowing us to coherently synthesize evidence from common garden and genome-environment associations. Our approach identifies loci exhibiting environmental clines where alleles are associated with higher fitness in home environments. Simulations show our approach can increase power to detect loci causing local adaptation. In a case study on *Arabidopsis thaliana*, most identified SNPs exhibited home allele advantage and fitness tradeoffs along climate gradients, suggesting selective gradients can maintain allelic clines. SNPs exhibiting G×E associations with fitness were enriched in genic regions, putative partial selective sweeps, and associations with an adaptive phenotype (flowering time plasticity). We discuss extensions for situations where only adaptive phenotypes other than fitness are available. Many types of data may point toward the loci underlying G×E and local adaptation; coherent models of diverse data provide a principled basis for synthesis.

## Introduction

Populations commonly exhibit phenotypic differences, often due to local adaptation to environment (Leimu & Fischer 2008; Hereford 2009). Local adaptation is defined as a genotype-by-environment interaction (G×E) for fitness that favors home genotypes (Kawecki & Ebert 2004). Local adaptation has long interested empirical and theoretical biologists (Clausen *et al.* 1940, 1948; Levene 1953; Slatkin 1973). However, little is known about the genomic basis of local adaptation, such as genetic architecture, major molecular mechanisms, and the extent to which genomic divergence among populations is driven by local adaptation. Because local adaptation involves organismal responses to environmental gradients, understanding the mechanisms of local adaptation has important applications in agriculture and biodiversity conservation (Aitken & Whitlock 2013; van Oppen *et al.* 2015; Lasky *et al.* 2015). Additionally, G×E are important in human phenotypes like disease (Anastasi 1958; Hunter 2005; Gage *et al.* 2016). Understanding the genomic basis of G×E is an emerging area of biomedical research (Thomas 2010; Keller 2014) as are the genomics of local adaptation (reviewed by Des Marais *et al.* 2013; Manel & Holderegger 2013; Tiffin & Ross-Ibarra 2014; Adrion *et al.* 2015; Bragg *et al.* 2015; Hoban *et al.* 2016).

A central question in local adaptation is whether selective gradients alone can maintain allelic clines at individual loci, or whether stochastic processes, like limited dispersal, are required to explain clines at individual loci causing local adaptation (Mitchell-Olds *et al.* 2007; Anderson *et al.* 2011b). If selective gradients cause rank changes in alleles with the highest relative fitness at an individual locus, selection may maintain a cline, a pattern known as genetic tradeoff or antagonistic pleiotropy (Ågren *et al.* 2013). Detecting loci that exhibit antagonistic pleiotropy in locally adapted systems has been challenging, partly due to limited statistical power of approaches that conduct multiple tests of significance for opposing fitness effects in different environments (Fournier-Level *et al.* 2011a; Anderson *et al.* 2013).

Common garden experiments have long been employed to characterize genetic variation in phenotypes (Langlet 1971). In particular, reciprocal common gardens at multiple positions along environmental gradients are a powerful tool to reveal local adaptation (Clausen *et al.* 1940, 1948). One approach to identifying the loci underlying local adaptation is to combine fitness data from multiple common garden experiments with genomic data (Lowry & Willis 2010; Fournier-Level *et al.* 2011a; Anderson *et al.* 2011a; Ågren *et al.* 2013). However, common gardens are logistically challenging and it is unclear how the typically small spatiotemporal scales of common gardens relate to the scales of processes that generate local adaptation in the wild (Weigel & Nordborg 2015).

An alternative to discovering genetic and ecological mechanisms of local adaptation is to study changes in allele frequency along environmental gradients (Hedrick *et al.* 1976; Tiffin & Ross-Ibarra 2014; Adrion *et al.* 2015; Bragg *et al.* 2015; Rellstab *et al.* 2015; Hoban *et al.* 2016). In this approach, known as a genome-environment association study, individuals are sequenced from multiple locations along environmental gradients. Genetic markers and environmental gradients showing the strongest correlations are then considered as potentially involved in local adaptation (*e.g.* Hancock *et al.* 2008, 2011; Eckert *et al.* 2010; Turner *et al.* 2010; Coop *et al.* 2010; Lasky *et al.* 2012; Jones *et al.* 2012; Fitzpatrick and Keller 2015). A challenge of both traditional association studies (genome-phenotype) and genome-environment association studies is that the genomic variation is observational and is not experimentally randomized (as opposed to linkage mapping with experimental crosses, Devlin & Roeder 1999; Hancock *et al.* 2008; Kang *et al.* 2008; Nordborg & Weigel 2008). As a result, many loci may show spurious associations with phenotypes or with environment (Price *et al.* 2010; Schoville *et al.* 2012; Bragg *et al.* 2015). Spurious associations are particularly problematic for environmental gradients that are spatially autocorrelated due to confounding with population structure (Schaffer & Johnson 1974). A technique for dealing with this confounding is to control for putative population structure when testing associations (Coop *et al.* 2010) by controlling for genome-wide relatedness (*e.g*. estimated by identity-in-state) among accessions (Yoder *et al.* 2014; Lasky *et al.* 2014).

Understanding the genomic basis of adaptation may benefit from synthesizing lines of evidence, for example by combining multiple types of genome scans to strengthen the evidence that a locus is under selection (Lasky *et al.* 2014; Evans *et al.* 2014). For example, researchers have identified overlap between outliers for selection statistics and markers associated with putatively adaptive phenotypes (Horton *et al.* 2012) or between SNPs associated with phenotypes and those associated with climate gradients (Berg & Coop 2014). Lasky *et al.* (2015) used a Bayesian approach to combine associations with phenotype and environment, first calculating climate associations and then using each marker’s association to determine the prior probability it was associated with G×E for adaptive phenotypes, yielding a posterior. Although combining multiple lines of evidence is potentially useful, the quantitative approaches in past studies have often been *ad hoc* and lacked reasoned principles. Here we develop a modeling framework to conduct genome-wide association scans for G×E while coherently synthesizing multiple data types. Existing approaches to genome-wide association studies (GWAS) with G×E (sometimes referred to as genome-wide interaction studies, GWIS) have dealt with categorical nominal environments (Murcray *et al.* 2009; Thomas 2010; Korte *et al.* 2012; Gauderman *et al.* 2013; Marigorta & Gibson 2014), benefiting from the statistical convenience of modeling phenotypes in different environments as correlated traits (Falconer 1952). Association models have not been applied to G×E along continuous environmental gradients, such as modeling SNP associations with reaction norms (Jarquín *et al.* 2014; Tiezzi *et al.* 2017). Despite the existence of studies where fitness was measured in multiple common gardens for diverse genotyped accessions (Fournier-Level *et al.* 2011a), studies where linkage mapping was conducted for fitness at multiple sites (Ågren *et al.* 2013), and studies where authors conducted association mapping for G×E effects on phenotypes (Li *et al.* 2014), we found no example of association studies of G×E for fitness, which is the basis of local adaptation.

The underlying processes generating local adaptation are the same regardless of the approach used for inference, be it genome-environment association or common gardens. Thus it is natural to synthesize data from multiple approaches. Furthermore, by combining data types into a single inferential framework we may increase our power to find causal loci. Here, we simultaneously leverage data from multiple common gardens and genome-environment associations by using a new approach to imputing relative fitness when common gardens are missing (Figure 1). In the remainder, we describe our approach, present test cases using simulations and published data on *Arabidopsis thaliana* (hereafter Arabidopsis), and discuss extensions.

**Figure 1.**
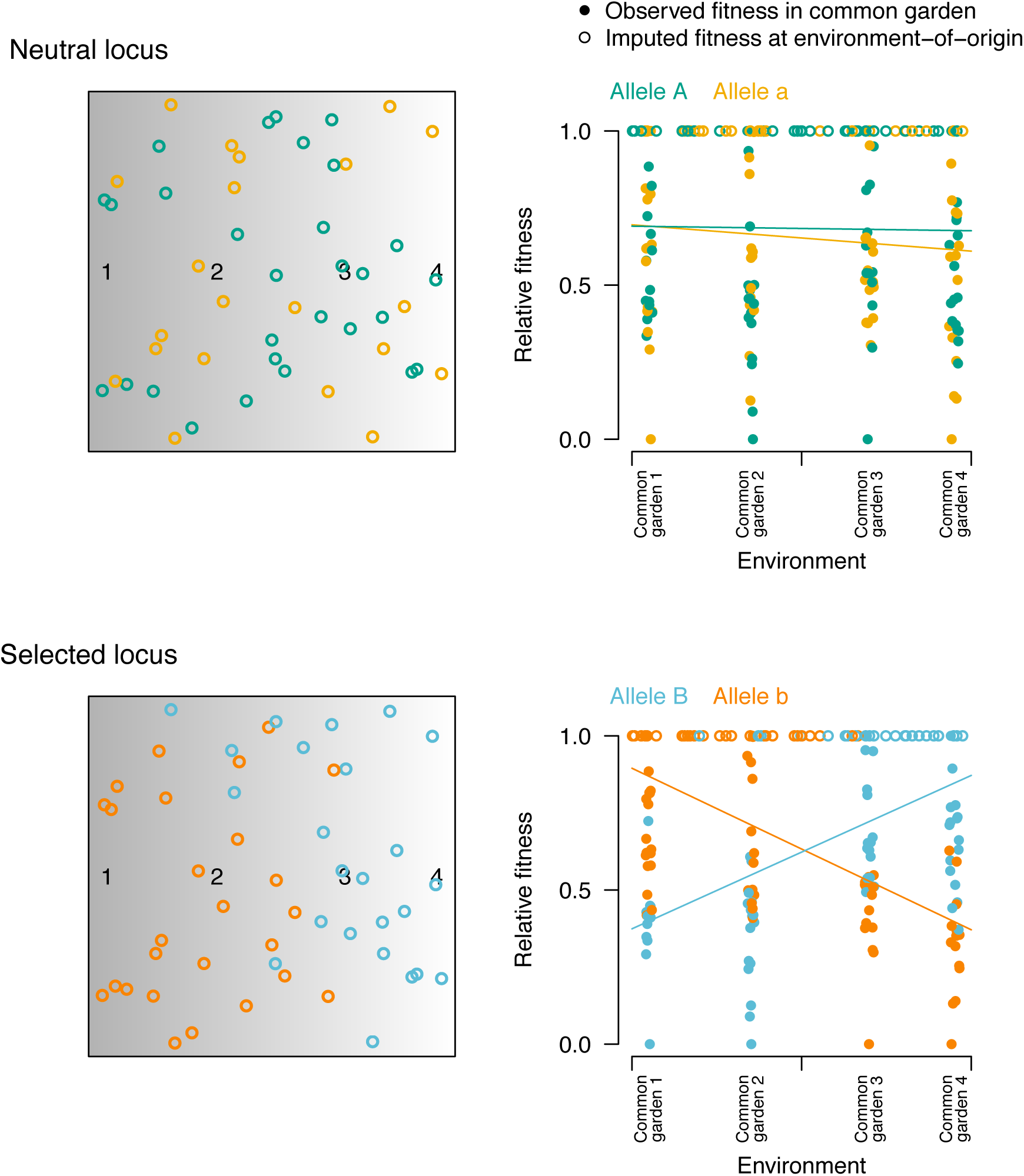
Illustration of our imputation technique and stereotypical patterns captured by our approach for neutral (top panels) and selected (bottom panels) loci. Here we show hypothetical data from four common gardens along an environmental gradient (solid circles in four vertical streaks in right panels, with small amount of noise added to environmental values for visualization) that have fitness scaled to a maximum of 1. We also show accessions (or ecotypes) collected in home environments and sequenced (mapped in two-dimensional geographical space in left panels) having imputed relative fitness of 1 in their home environment (environment-of-origin, open circles at top of right panels). The selected locus (bottom right panel) in question shows strong G×E for fitness, such that allele *B* (blue) is more fit (observed, solid circles) and more common (imputed, open circles) at the right side of the environmental gradient while allele *b* (orange) is more fit and more common at the left side of the gradient.

## Methods

### Genome-wide association study of G×E effects on fitness

In common garden experiments, environment is often treated as a factor. When more than two gardens are conducted, variation among them may be considered in a more general fashion. For a given environmental gradient, each common garden may be located along the gradient according to its conditions. Describing common gardens as such may be informative about the specific ecological mechanisms driving selective gradients, taking advantage of the ordered nature of gardens’ environments. We leverage multiple common garden experiments to identify markers (single nucleotide polymorphisms, SNPs) that show the strongest G×E effects, loci where allelic state shows the strongest interaction with environment in its association with fitness.

Local adaptation requires a genotype by environment interaction for fitness at the whole genome-level. Variation in individual phenotypes from multiple environments can be separated into components determined by genotype, environment, and G×E (Yates & Cochran 1938; Falconer 1952). To assess this interaction at an individual locus, one can assume that the relative fitness of individual *i* in a single location, *w*_*i*_, satisfies the linear model

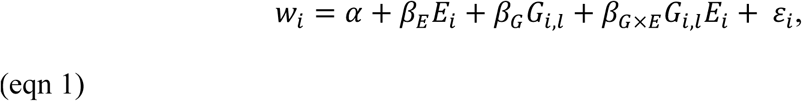

where *G*_*i,l*_ is the genotype of individual *i* at locus *l* and *E*_*i*_ is the value of a single environmental variable at the location where *w*_*i*_ was measured. The *β*_E_ parameter gives the effect of environment on fitness and *β*_G_ gives the effect of genotype on fitness. Our primary interest lies in the *β*_G×E_ parameter, which gives the strength and direction of G×E effects; *β*_G×E_ determines how responses to environmental gradients are mediated by genotype. The term *α* gives the fitness intercept. We assume that the vector of errors, ***ε***, can be expressed as

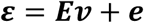

where ***E*** is a diagonal matrix of the environmental values, and

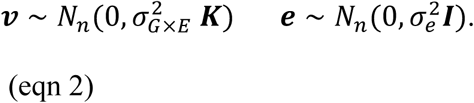

Here ***v*** and ***e*** are independent. The *n*×*n* matrix **K** is calculated as the genome-wide identity in state for each pair of the sampled *n* accessions (Kang *et al.* 2008). Random effects ***v*** are included because a substantial portion of G×E may be associated with population structure (Lasky *et al.* 2015); naively applying standard F-tests to assess the interaction effects can result in a dramatic increase in false positive rates. To ameliorate this issue, the random effect ***v*** represents the genetic background interactions with environment (G×E, magnitude of their variance determined by 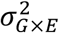), while ***e*** represents the independent and identically distributed error in the model (variance determined by 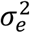:). However, incorporating random effects may also decrease power when causal loci covary with genomic background.

### Coherent synthesis of common gardens and genome-environment associations via imputation

We now tackle the goal of synthesizing genome-environment associations and G×E observed across common gardens. A challenge in synthesizing these approaches is that genome-environment studies are purely observational and lack common garden experiments. However, an implicit assumption in studies of genome-environment associations is that local adaptation occurs; if a common garden were conducted at each location where wild genotypes are collected, the home genotype would tend to be most fit. Our imputation approach makes this assumption explicit. A formal consequence of this assumption is an (imputed) observation of highest relative fitness for genotypes at home. We combine this imputation with two observed datasets: genomic markers and environment of origin (Figure 1). Next, we scale relative fitness within each common garden so that the maximum observed fitness is given a relative fitness of unity, yielding a metric that can be directly observed or imputed in each type of study (common garden or genome-environment association). For imputation, we then assume that each genotype collected in the wild is locally adapted at its home and thus has a relative fitness = 1 (Figure 1). After imputation, we can calculate marker associations with G×E for fitness, where each fitness observation arises from either (A) observations on a given genotype by common garden combination or (B) imputation on a given genotype collected from its natural home and subsequently sequenced.

### Fitting models and comparing approaches

We compared four reaction norm approaches to genome-wide G×E association studies in addition to the more common approach of genome-environment associations. In Approach 1, we ignored potential confounding of population structure, using least squares to fit the model in eqn 1 where ***ε*** is normal, independent, and identically distributed (excluding random effects ***v***), but only including observed fitness data from common gardens and excluding imputed fitness data. In Approach 2, we again used least squares but included imputed fitness data; these imputations using information from the ancillary geographic data could possibly reduce false positives. In Approach 3, we fit the full mixed-effects model (including random effects ***v***), but excluded imputed fitness. In Approach 4, we fit the full mixed-effects model while including imputed fitness data. To test a genome-environment association approach (Approach 5), we also compared associations between SNPs and home environments used a mixed-effects model in an approach akin to traditional association mapping but where environment is substituted for phenotype (Yoder *et al.* 2014; Lasky *et al.* 2014).

To improve computation time for the mixed-model approaches (3-5), we used the method of Kang *et al.* (2010) and first fit the random effects with covariance determined by kinship, and then fixed these effects while testing the effects of each SNP on the phenotype. We included the environmental covariate effect in this initial step, following the recommendation of Kang *et al.* (2010) for fitting additional non-SNP covariates. In other words, we first fit the model:

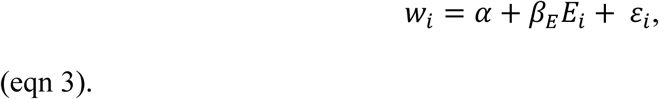

Eqn. 3 is the same as Eqn. 1 but with genetic effects omitted. We obtained parameter estimates 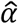, 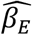, 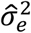, 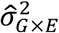. We then take the variance parameter estimates and use them to estimate the remaining slope coefficients in eqn 1 using generalized least squares. Because inclusion of the *β*_*E*_ term in ordinary least squares regression (Approaches 1-2) led to poor model fit, we excluded the term from those approaches. We fit the discussed mixed-model using Minimum Norm Quadratic Unbiased Estimation, MINQUE (Rao 1971; Brown 1976; Reimherr & Nicolae 2016). This approach is equivalent to restricted maximum likelihood, REML, but rephrased in a way that more fully exploits the linearity of the model, resulting in a flexible framework that can be quickly computed.

### Simulation

We used simulations to demonstrate how our imputation technique can improve power to identify loci causing G×E for fitness. To assess scenarios with varying strength of local adaptation, we tested simulations of varying dispersal distances. We used results of Forester *et al.* (2016), who previously simulated local adaptation along a continuous environmental gradient (Landguth & Cushman 2010). Specifically, we used genome and environment data from Forester *et al.* (2016) and here simulated new common garden data. Forester *et al.* (2016) simulated 100 bi-allelic loci, one of which was under selection (more detail is found in Forester *et al.* 2016 and the Supplement for this paper). In the simulation, selection changed linearly along an environmental gradient, with *AA* and *aa* genotypes favored at different ends of the gradient and *Aa* genotypes under uniform selection across the gradient (Fig. S1). We tested three values of dispersal parameters (low to high), resulting in varying strength of local adaptation, with the Pearson’s correlation between selected locus and selective gradient equal to 0.28, 0.24, and 0.11, respectively.

Here, we sampled 250 individuals randomly. We then located four common gardens at equal intervals along the gradient, encompassing the extremes of the selection surface (Fig. S1). For the moderate dispersal and local adaptation scenario, we tested the effect of common gardens that sample only half the environmental gradient (Fig. S1). For the gardens, we subsampled 100 individuals from the full 250, and then averaged fitness for 25 clones of each individual (each with the identical adaptive genotype of their parent clone) in each common garden using the selection model of Forester *et al.* (2016). After imputing fitness for the 250 individuals in their home environments, we had a total of 650 observations of fitness × location (250 imputed observations from individuals sampled across the landscape + 400 real observations arising from 100 clones in each of four common gardens).

For both the simulations and the *Arabidopsis* case study, we focus on the 1% of SNPs with the lowest p-values and their role in local adaptation. In simulations with 100 SNPs, this was equivalent to the lowest p-value SNP. To determine false positive rates in simulations, we calculated the proportion of simulations for a given scenario where a neutral (as opposed to a causal) SNP had the lowest p-value for *β*_G×E_.

### Model extensions

Two extensions to our approach could increase its generality. First, one could treat unobserved fitness of a genotype in its home environment as a free parameter. To constrain estimates of unobserved fitness one could use informative priors, such that the prior probability of relative fitness at home for each genotype would be monotonically increasing, *i.e.* local adaptation is the most likely state, but minor maladaptation is common. Inferences about unobserved fitness could be further constrained using hierarchical models, such that home fitness parameters for multiple genotypes arise from a common distribution (Gelman & Hill 2007). Relaxing the assumption of perfect local adaptation would also generate less biased, if less precise, parameter estimates for *β*_G×E_, which are currently conservative when imputation of local adaptation for maladapted genotypes pushes *β*_G×E_ toward zero and weakens estimates for selective gradients.

Second, when fitness is not measured, components of fitness (*e.g.* survival) or traits thought to be locally adaptive (*e.g*. physiological) can be measured and used to infer the genomic basis of local adaptation. For example, instead of modeling SNP×environment associations with fitness, one could model SNP×environment associations with components of fitness measured in common gardens, and estimate unobserved traits for sequenced genotypes using informative priors. To be clear, in our case study of Arabidopsis, we had near but not complete lifetime fitness data (missing germination stage). Here we do not fit these model extensions to data, given the current computational challenge of fitting many more parameters in a Bayesian framework.

### *Case study: local adaptation to climate in* Arabidopsis thaliana

We applied these approaches to published data from studies of *Arabidopsis thaliana* in its native Eurasian range. Fournier-Level *et al.* (2011a) conducted replicated common gardens at four sites across Europe: Spain, England, Germany, and Finland (Figure 2). With these data, Fournier-Level *et al.* (2011a) and Wilczek *et al.* (2014) showed evidence that genotypes are locally adapted to their home temperature and moisture regimes and that alleles associated with high fitness in a given garden tended to be found nearer to that garden than alternate alleles, suggesting these loci were involved in local adaptation. At each site the authors transplanted 157 accessions (59 in the case of Finland) on a date in the fall matched to germination of local winter-annual natural populations (Fournier-Level *et al.* 2011a). The authors calculated survival (out of individuals surviving transplant) and average fecundity (where individuals that died before reproducing had fecundity zero) giving an estimate of absolute fitness (excluding the seed to seedling transition, Fournier-Level *et al.* 2011a).

**Figure 2.**
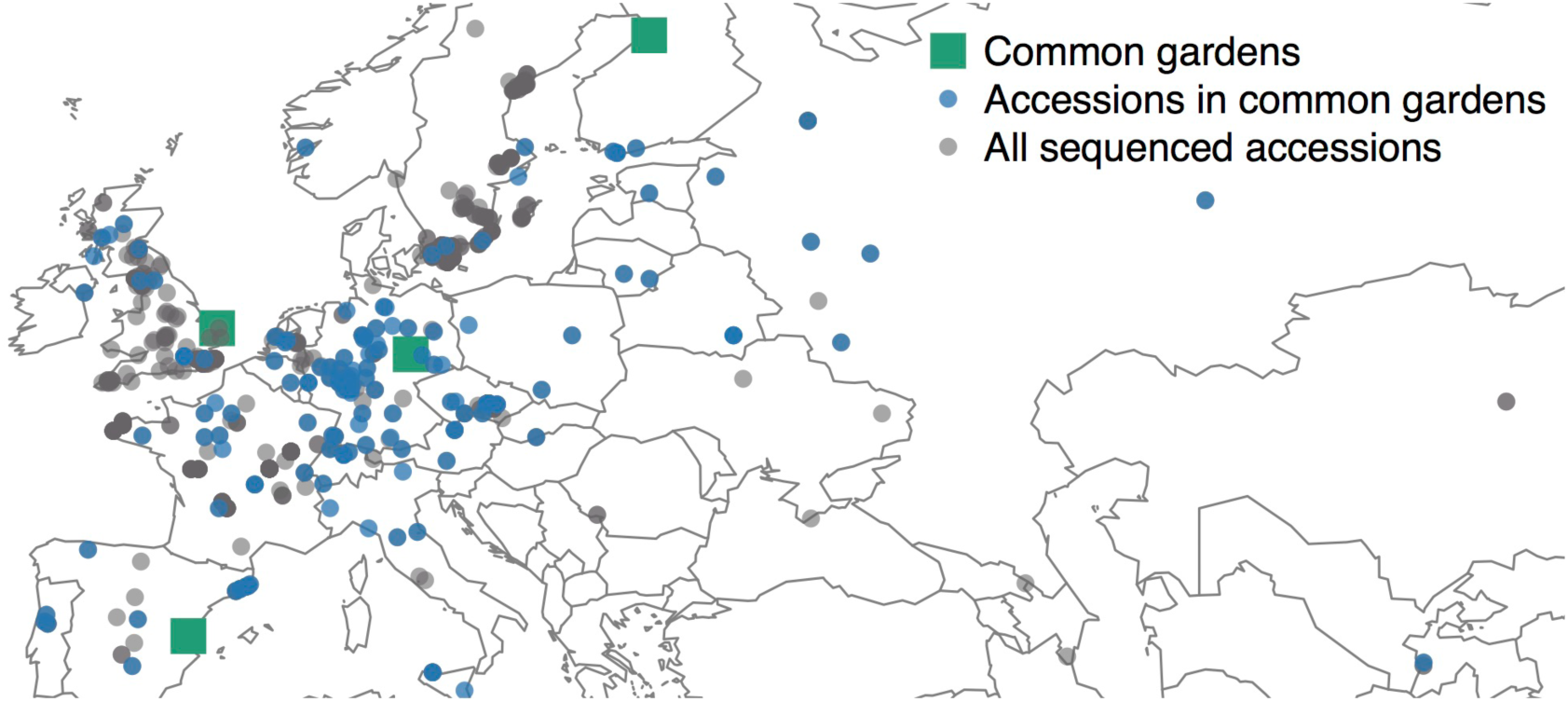
Data used in case study on Arabidopsis. The location of common gardens, natural accessions in common gardens, and all other sequenced natural accessions are shown.

These accessions were part of a panel of 1,307 accessions from around the globe that were genotyped at ∼250k SNPs using a custom Affymetrix SNP tiling array (AtSNPtile1), with 214,051 SNPs remaining after quality control (Figure 2, Horton *et al.* 2012). This array was generated by resequencing 19 diverse ecotypes from across the range of Arabidopsis (Kim *et al.* 2007). Of the 1,307 genotyped accessions, we used 1,001 accessions that were georeferenced and likely not contaminant lines (Anastasio *et al.* 2011), in addition to being from the native range in Eurasia (Hoffmann 2002; Lasky *et al.* 2012b), and excluding potentially inaccurate high altitude outliers (i.e. > 2000 m). After imputing fitness for accessions in their home environments we had a total of 1,531 observations of fitness × location (1001 imputed observations + 530 real observations arising from the four common gardens). We removed from association tests SNPs having minor allele frequency (MAF) below 0.01, though we also considered a more conservative threshold of MAF = 0.1.

We used climate and derived data from a previous study (Lasky *et al.* 2012b) using published global climate datasets (Hijmans *et al.* 2005; Zomer *et al.* 2008). Here we focus on four climate variables that differ among common gardens, are not strongly correlated, and may be involved in local adaptation: minimum temperature of the coldest month, average monthly minimum temperature in the growing season, coefficient of variation of monthly growing season precipitation, and aridity index.

### Genome-wide G×E associations

We separately tested for each SNP’s interaction with each of the four climate variables. For each of the four approaches using common garden data we fit a model for each combination of SNP and environmental variable. To characterize the types of patterns identified by our approach, we studied variation in the SNPs in the 0.01 lower tail of p-values for the hypothesis test that *β*_G×E_ = 0 (the coefficient for SNP*×*environment effects on fitness) for each climate gradient. For these SNPs, we calculated whether the direction of allelic differentiation along environmental gradients was consistent with the sign of *β*_G×E_. For example, if one allele was more common in accessions from warmer locations, we assessed whether that same allele showed an increase in relative fitness in warmer common gardens.

Next, we assessed whether our model predicted that different alleles were most fit in the two common gardens at either extreme of a climate gradient, *i.e.* whether the SNP was associated with rank changes in fitness that are consistent with genetic tradeoffs, versus a pattern where *β*_G×E_ merely involved changes in fitness difference between alleles (such as conditional neutrality or variance changing G×E), the latter of which cannot stably maintain local adaptation. For example, if one allele was estimated to be most fit in the coldest common garden, we determined whether a different allele was estimated to be most fit in the warmest common garden. Furthermore, we also quantified similarity (rank correlation in SNP scores and proportion of SNPs common to the strongest 0.01 tail of associations) between results from our G×E approach versus those from other recent studies of association with home climate in Arabidopsis (Hancock *et al.* 2011; Lasky *et al.* 2012b, 2014).

### Enrichment of strong SNP×environment associations across the genome

We studied whether loci we identified as likely being involved in local adaptation exhibited supportive patterns in ancillary datasets. First, to assess whether our association approach is capable of identifying the signal of local adaptation rather than spurious background associations, we tested for enrichment of SNPs in genic (from transcription start to stop sites) versus intergenic regions. These tests are based on the hypothesis that loci involved in adaptation are on average more likely to be found near genes and linked to genic variation, in comparison with loci evolving neutrally (Hancock *et al.* 2011; Lasky *et al.* 2012b). For a test statistic, we calculated the portion of SNPs in the 0.01 lower p-value tail that were genic versus intergenic.

Second, we hypothesized that locally-adaptive alleles may have been subject to partial (local) selective sweeps, especially given that much of Arabidopsis’ Eurasian range was recently colonized following the last glacial maximum. We tested for an enrichment of significant (alpha = 0.05) pairwise haplotype sharing (PHS, Toomajian *et al.* 2006) in the SNPs (using PHS calculated by Horton et al. 2012) showing the greatest evidence of G×E for fitness. We also tested evidence that these SNPs are enriched for significant (alpha = 0.05) integrated extended haplotype homozygosity (standardized, iHS, Voight *et al.* 2006), an additional metric of partial sweeps. We used ancestral SNP allele determinations from Horton *et al.* (2012, based on alignment with the *Arabidopsis lyrata* genome) and the R package ‘rehh’ to calculate iHS (Gautier *et al.* 2012).

Third, we also studied whether loci we identified were associated with plasticity in flowering time, a trait that plays a major role in local adaptation to climate in plants (Hall & Willis 2006; Franks *et al.* 2007; Keller *et al.* 2012; Lowry *et al.* 2014). Recently Li *et al.* (2014) tested the flowering time response of 417 natural accessions to simulated warming (up to ∼4ºC), and then identified SNP associations with changes in flowering time across treatments, G×E for flowering time. We tested whether SNPs we identified as having SNP×environment interactions for fitness (0.01 lower p-value tail) were enriched in nominally significant associations (alpha = 0.05) with G×E for flowering time.

To generate a null expectation for each enrichment while maintaining a signal of linkage disequilibrium in the null model, we circularly permuted SNP categories (*e.g*. as genic versus intergenic, having significant iHS or not) along the genome and recalculated the test statistics 10,000 times (Hancock *et al.* 2011; Lasky *et al.* 2012b).

## Results

We compared four approaches to genome-wide G×E association study and one approach for genotype-environment association. Approach 1 used only observed (excluding imputed) fitness data, but no correction for population structure. Approach 2 used observed and imputed fitness data, and no correction for population structure. Approach 3 fitted the full mixed-effects model, but only including observed fitness data from common gardens, excluding imputed fitness. Approach 4 fitted the full mixed-effects model while including both observed and imputed fitness data. Approach 5 used a mixed model of genotype associations with environment (no common garden data).

### Simulations

Across dispersal scenarios, we found that mixed models decreased false positive rates and increased accuracy of inference as to the SNPs driving G×E for fitness (Figure 3). When dispersal was highest and local adaptation weakest, all approaches exhibited an increase in false positives compared to the moderate dispersal scenario. Among the approaches using common garden data (Approaches 1-4), the mixed models (Approaches 3-4) generally had low false positive rates and thus high true positive rates compared to models excluding random effects (Approaches 1-2). Based on the low false positive rates and low p-values for causal SNPs in Approach 3, common garden data were a clear source of statistical power to identify causal SNPs (Figure 3). Including imputed data (Approach 4) further reduced false positive rates and resulted in no false positives under the two lower dispersal scenarios. Genotype-environment associations that did not use common garden data (Approach 5) had similarly low false positive rates (Table S5). However, common garden data combined with imputations (Approach 4) yielded stronger inference for SNPs driving G×E; causal SNPs had lower p-values (Figure 3, median causal SNP p for low, medium, high dispersal scenarios: 3.2*×*10^-6^, 4.5*×*10^-13^, 7.4*×*10^-7^) compared to ignoring common garden data (Approach 5, median causal SNP p for low, med., high dispersal scenarios: 2.3*×*10^-7^, 1.9*×*10^-10^, 1.2*×*10^-5^). Under a scenario of medium dispersal and common gardens that only covered half the gradient, false positive rates were elevated for approaches excluding random effects (Approaches 1-2) or excluding imputations (Approach 3) but not when imputations and random effects were included (Approach 4, Figure S4).

**Figure 3.**
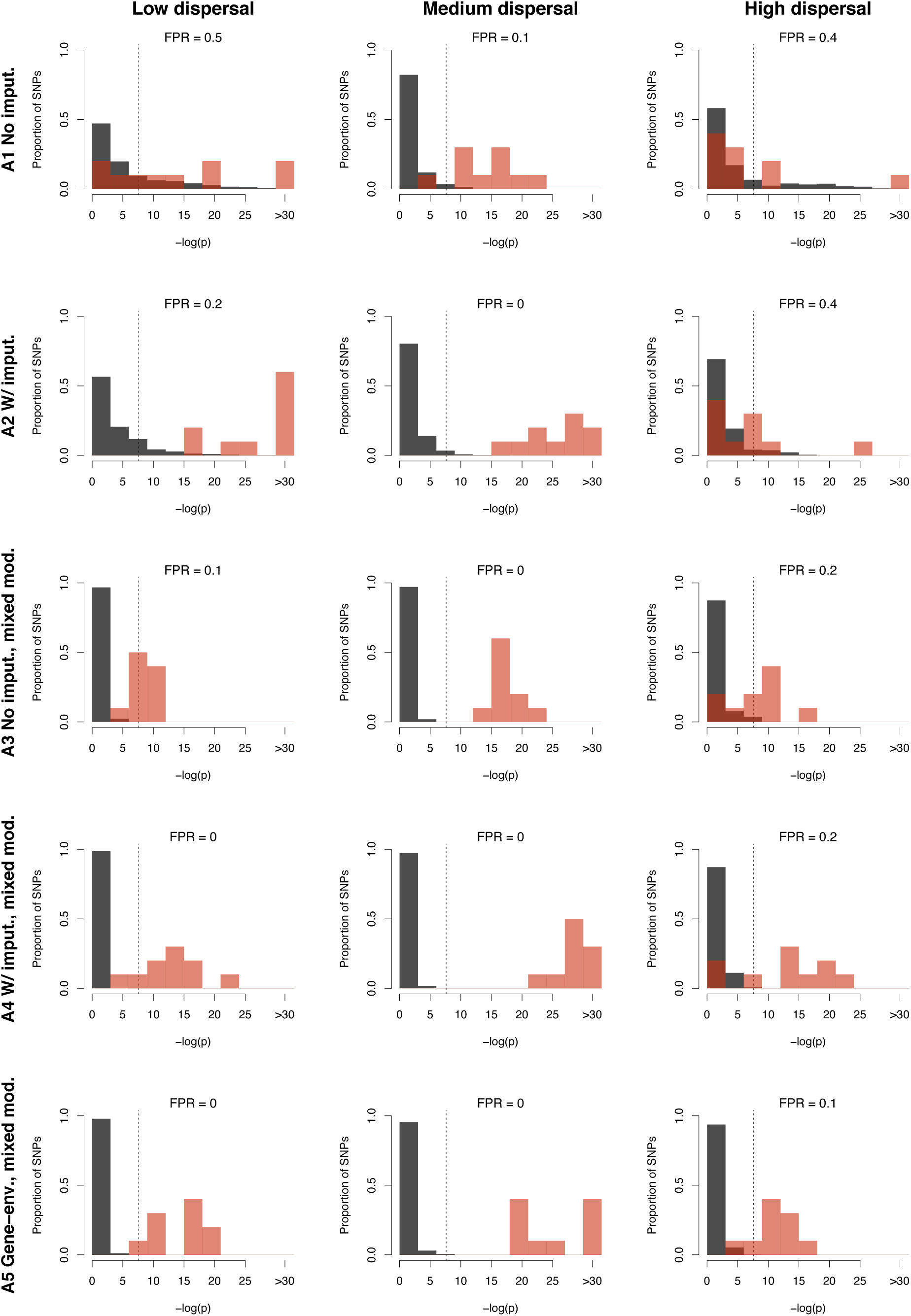
Comparison of causal (red) and neutral (black) SNP associations with G×E for fitness across three different levels of dispersal and 10 replicate simulations for each level. Approaches used (row 1) no imputation and no random effects, (row 2) imputation but no random effects, (row 3) mixed models that used only observations from four common gardens, (row 4) mixed models combining imputed observations of relative fitness in home environments with common garden observations, or (row 5) a genotype-environment association approach. False positive rate (FPR) is indicated, calculated as the proportion of simulations where a neutral SNP had the lowest p-value. Each simulation had 1 causal SNP and 99 neutral SNPs; plots show aggregate distributions for all SNP by simulation combinations (*i.e.* total of 10 causal and 990 neutral SNPs). Y-axes show the proportion of SNPs in each category (causal or neutral) falling into a given p-value bin. For reference, dashed line indicates a strict Bonferroni cutoff for alpha = 0.05, –log(0.05/100) = 7.6.

### Case study on Arabidopsis

We found that simple linear model tests (Approaches 1-2) of SNP*×*environment interactions were highly enriched in very low p-values (Figure S5) relative to the theoretical expectation. After incorporating the kinship*×*environment random effects (but excluding imputed fitness observations, Approach 3), we found that SNP*×*environment associations with fitness were closer to the theoretical expectation but still highly enriched in low p-values for three climate variables. After incorporating imputed fitness observations into the mixed model (Approach 4, right column, Figure S5), we found p-value distributions hewed closer to the theoretical expectation and were slightly conservative (under-enriched in low p-values) for two climate variables. These approaches tended to identify different SNPs as having the strongest SNP*×*environment associations with fitness (Table S6).

Based on the results of our simulations and the p-value distributions noted above, we focus the remainder of analyses on results from mixed models with imputed fitness included (Approach 4). We found that climate variables differed in the importance of kinship-climate interaction associations with fitness (proportion of variance in fitness explained by random effects ***v***), suggesting that population structure in Arabidopsis is more strongly correlated with some climatic axes of local adaptation (G×E for fitness) compared to other climate gradients. For growing season minimum temperatures, kinship×environment interactions explained most of the variation in fitness (R^2^ =0.78, Table 1, row 3). By contrast, kinship×environment interactions for fitness were weaker along a gradient in winter minimum temperature (R^2^ =0.07).

**Table 1.** Characterization of patterns in *Arabidopsis* case study identified by Approach 4 (mixed model including imputations) for SNPs in 0.01 lower tail of p-values for SNP*×*environment interactions for fitness (first two rows of table) and for kinship*×*environment interactions for fitness (third row). SNPs with home allele advantage are defined as those where the sign of allelic differences in home climates were matched by the sign of fitted mixed model SNP*×*environment associations with relative fitness. Rank changing SNPs are those where we estimated a rank change in relative fitness for alternate alleles along the environmental gradient between the two extreme common gardens. The final row gives the proportion of total observed climate gradient (among ecotypes) captured by the two most extreme common gardens.

| *Statistic* | *Aridity* | *CV grow.*<br>*seas. prec.* | *Min. temp.* | *Min. temp.*<br>*grow. seas.* |
| --- | --- | --- | --- | --- |
| Proportion SNP×E with home allele advantage | 0.46 | 0.89 | 0.92 | >0.99 |
| Proportion SNP×E rank changing | 0.31 | 0.92 | >0.99 | 0.73 |
| Kinship×E $R^2$ for fitness | 0.08 | 0.58 | 0.07 | 0.43 |
| Proportion climate gradient covered by gardens | 0.13 | 0.65 | 0.78 | 0.31 |

Approach 4 tended to identify SNPs where SNP*×*environment interactions favored alleles in climates where they were relatively more common, that is, the sign of allelic differences in home climates were matched by the sign of fitted mixed model SNP*×*environment associations with relative fitness (Table 1, row 1, and see outlier examples in Figure 4). In addition to characterizing SNP*×*environment associations, we tended to identify SNPs where we estimated a rank change in relative fitness for alternate alleles along the environmental gradient between the two extreme common gardens (where the fitted model expectation was that the allele with higher fitness at one extreme common garden differed from the allele with higher fitness at the other extreme, Table 1, row 2). It appeared that the proportion of SNPs expected to show rank changes in relative fitness among the common gardens was related to how much of each climate variable’s range was covered by gardens (Table 1, row 4). Thus the common gardens may have been limited in their ability to capture rank changing of alleles at some loci involved in local adaptation to aridity and growing season cold.

**Figure 4.**
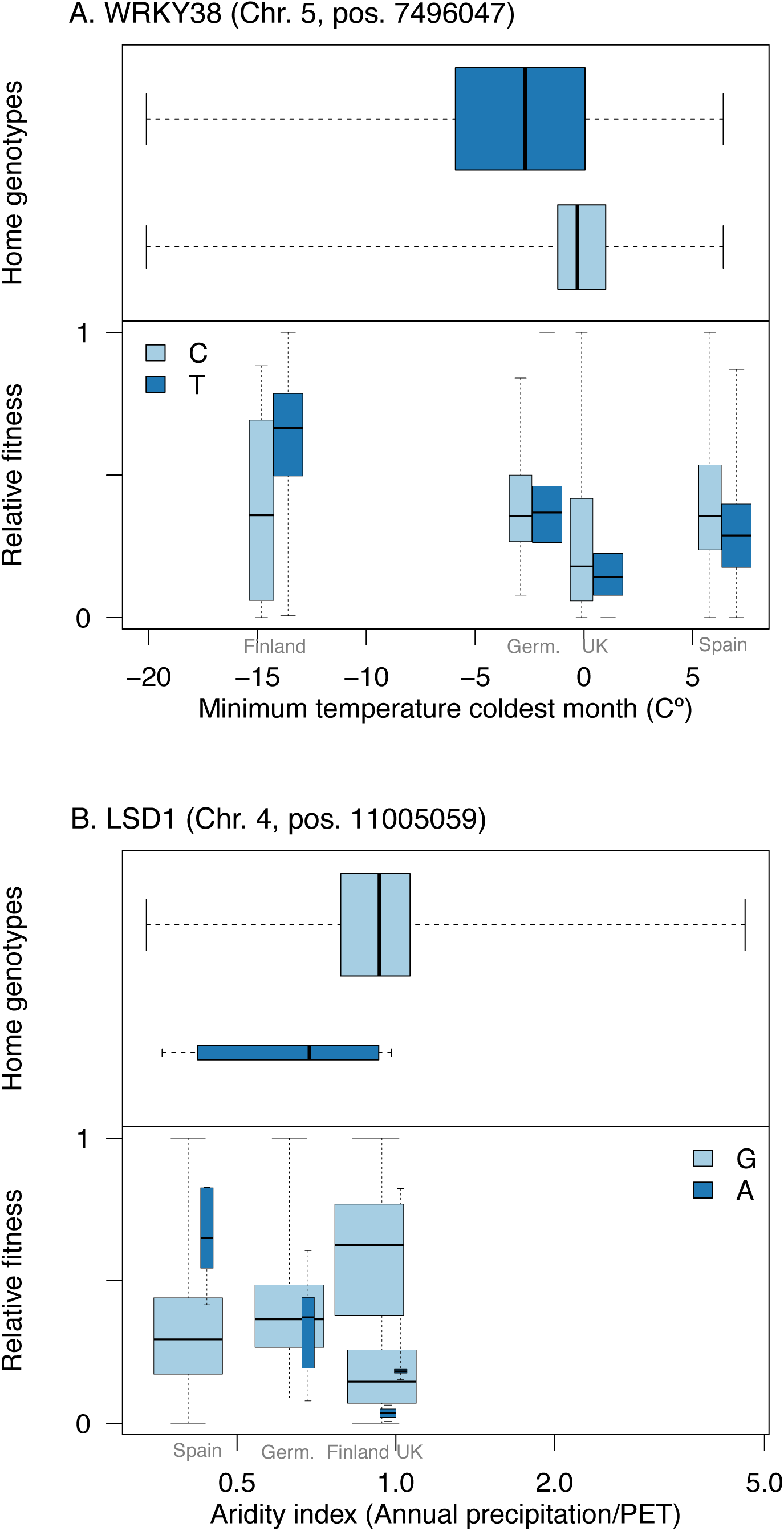
Example SNPs with the strongest associations (lowest p-values) with cold winter temperatures (A) and aridity (B). Top subpanels show the climate distribution of alleles in home genotypes (natural geographic patterns), known as genotype-environment associations. Bottom subpanels show relative fitness of alleles in four common gardens, where common gardens’ climates determine position on x-axes. Each SNP falls within the coding region of indicated genes (WRKY38 and LSD1). Box widths are scaled to relative number of accessions having each allele. In both (A) and (B), the allele with the greatest relative fitness in common gardens changes along the environmental gradient consistent with change in allele frequency in native accessions (*i.e*. ecotypes).

We found non-random, but very weak overlap between the SNPs we identified and those outliers in previous analyses (Hancock *et al.* 2011; Lasky *et al.* 2012b, 2014). When considering mixed model (Lasky *et al.* 2014) or partial Mantel (Hancock *et al.* 2011) SNP associations with the same climate variables (genome-environment associations with no common garden data), we found significant overlap among the previously identified SNPs in the 0.01 lower tail of p-values versus those in the 0.01 tail identified here (permutation test, all p<0.05, Table S7). However, rank correlations among SNP scores from previous approaches versus our current approach were very weak (all rho < 0.2, Table S7, Figure S6).

### SNP×environment associations with fitness are enriched in regions suggestive of local adaptation

We found that SNP*×*environment interactions for fitness were significantly enriched in genic regions (again focusing on Approach 4: mixed model including imputations, Table 2; for reference, SNPs identified via Approach 2, including imputation but without random effects, were not significantly enriched in genic regions). Additionally, we found that SNP*×*environment interactions for fitness were enriched for high pairwise haplotype sharing (PHS) and high integrated extended haplotype homozygosity (iHS, Table 2). Finally, SNPs associated with G×E for flowering time response to growing temperature (Li *et al.* 2014) tended to also have strong SNP*×*growing season minimum temperature interactions for fitness (p < 0.0002) but not for other climate variables. (Table 2) Enrichments reported above did not change qualitatively (with respect to statistical significance) when we only considered SNPs with MAF > 0.1.

**Table 2.** Permutation tests of enrichment p-values (Approach 4) for various signals suggestive of local adaptation to climate in case study on Arabidopsis. For each statistic, we tested for enrichment of signal in the SNPs in the 0.01 lower tail of p-values for SNP*×*environment associations with relative fitness. “Genic” tests enrichment of genic versus non-genic SNPs, “PHS” and “iHS” test for enrichment with significant (alpha = 0.05) pairwise haplotype sharing and standardized integrated extended haplotype homozygosity, respectively. The final row shows enrichment with SNPs having significant (alpha = 0.05) associations with change in flowering time (G×E) in response to warming during growth.

| *Statistic* | *Aridity*<br>*enrichment* | *CV grow.*<br>*seas. prec.*<br>*enrichment* | *Min. temp.*<br>*enrichment* | *Min. temp.*<br>*grow. seas.*<br>*enrichment* |
| --- | --- | --- | --- | --- |
| Genic | 0.0036 | <0.0002 | 0.0054 | 0.0006 |
| PHS | 0.0272 | 0.0122 | 0.0068 | 0.0272 |
| iHS | 0.2840 | 0.0006 | 0.0020 | 0.0008 |
| Flowering time under<br>warming, G×E | 0.8582 | 0.0680 | 0.5844 | <0.0002 |

### SNP×environment associations with fitness identify genes potentially involved in local adaptation

Our approach identified a number of strong candidates for local adaptation at the top of lists of SNPs with the strongest SNP*×*environment associations with relative fitness (Tables S1-S4). For example, the top SNP associated with aridity interaction effects on fitness (chr. 4, position 11005059) fell within LESION SIMULATING DISEASE 1 (LSD1), which affects a number of traits in Arabidopsis, including survival and fecundity under drought (Wituszyńska *et al.* 2013; Szechyńska-Hebda *et al.* 2016, Figure 4), while the third SNP (chr. 2, position 7592008) fell within ATMLO8, MILDEW RESISTANCE LOCUS O 8, homologous with barley MLO which controls resistance to the fungal pathogen powdery mildew (Büschges *et al.* 1997). The top SNP associated with winter cold interaction effects on fitness (chr. 5, position 7496047) falls within coding region of WRKY38, involved in the salicylic acid pathway and pathogen defense (Kim *et al.* 2008), and was the same locus identified as most strongly associated with multivariate climate in Lasky et al. (2012, Figure 4). The top SNP associated with variability in growing season precipitation interaction effects on fitness (chr. 2, position 18504858) falls 380 bp from ABA HYPERSENSITIVE GERMINATION 11, AHG11, which mediates the effect of abscisic acid (ABA), a major hormone of abiotic stress response, on germination (Murayama *et al.* 2012). The fifth highest SNP (and second highest locus) associated with growing season cold interaction effects on fitness (chr. 3, position 8454439) fell within ABERRANT LATERAL ROOT FORMATION 5, ALF5, a gene that confers resistance to toxins (Diener *et al.* 2001) belonging to the MATE gene family, which play a variety of roles responding to environment (Shoji 2014).

## Discussion

Genetic variation in environmental responses (G×E) is ubiquitous but its genetic and physiological basis and role in local adaptation is poorly understood. Replicated common garden experiments and genome scans for loci exhibiting evidence for local adaptation have been important in understanding the genetic basis of G×E and local adaptation (Hancock *et al.* 2008; Eckert *et al.* 2010; Turner *et al.* 2010; Fournier-Level *et al.* 2011a; Lasky *et al.* 2012b; Ågren *et al.* 2013; Evans *et al.* 2014; Lasky *et al.* 2015). However, the complementary information in common gardens and geographic variation in allele frequency have not been coherently synthesized. Previous association studies of G×E have modeled discrete, categorical environmental effects (Murcray *et al.* 2009; Thomas 2010; Korte *et al.* 2012; Marigorta & Gibson 2014). The modeling of G×E across discrete, categorical environments is often employed for mathematical convenience, as such a treatment allows the use of models designed for multiple phenotypes, where the same phenotype in different environments is considered as multiple phenotypes (Falconer 1952).

We demonstrated an approach to association study of G×E for fitness and an imputation technique that allowed us to coherently synthesize evidence from common gardens and genome-environment associations. Our imputation method relied on making explicit the implicit assumption of local adaptation that underlies genome-environment association studies (Coop *et al.* 2010; Hancock *et al.* 2011; Lasky *et al.* 2012b). Using simulation, we demonstrated that this imputation can increase power to identify SNPs causing G×E for fitness and local adaptation. Our approach also identified strong candidate genes in Arabidopsis associated with SNPs that exhibit fitness tradeoffs along climate gradients such that locally common alleles had greater relative fitness.

The relative information on selective and adaptive genetic mechanisms contained in the two datasets (common garden, geographical genomic) for a given system will be determined by several factors. First, the power of common gardens depends on the range of sampled covariates (genotype and environment). We found evidence with both our simulations and empirical case study that greater coverage of environmental gradients can increase power to detect causal loci. Similarly, power may be enhanced by including in common gardens a range of variation at locally adaptive loci using diverse germplasm from across gradients. However, alternate mechanisms of local adaptation across regions and confounding between population structure and adaptive loci suggest that regional stratification in scans for local adaptation may be more powerful (Horton *et al.* 2016). Additionally, the power of common gardens is influenced by the match between conditions in gardens and long-term natural selective gradients that give rise to local adaptation (Weigel & Nordborg 2015), and the heritability of adaptive traits and fitness. The information contained in genome-environment associations (and hence imputed fitness data here), is influenced by the strength of local adaptation in sampled populations (Figure 3), which itself is determined by steepness of selective gradients, the level of gene flow, and time populations have had to evolve toward equilibrium allele frequencies (Yeaman & Whitlock 2011; Lotterhos & Whitlock 2014; Forester *et al.* 2016). It is important to recognize that our simulations covered a limited range of the parameter space relevant in nature (genetic architecture of local adaptation, dimensionality of environmental selective gradients, *etc*.). Here, populations were given time to reach equilibrium (Forester *et al*. 2016), which likely enhanced the power of genotype-environment associations compared to scenarios common in nature where populations are still responding to environmental changes. Apart from information on genetic mechanisms of G×E for fitness, common gardens afford a more direct opportunity to study phenotypes under selection, as opposed to genotype–environment associations where information on phenotype is limited to gene annotations.

Above we described a method of imputation based on the assumption of local adaptation, *i.e*. home genotypes had greater fitness than away genotypes. However, local adaptation in nature is typically imperfect, such that the optimal genotype for a given location might not be the home genotype (Leimu & Fischer 2008; Hereford 2009). Local adaptation may not occur for several reasons, such as gene flow across environmental gradients (Slatkin 1973), limited genetic variation (Barton 2001), temporal environmental shifts, and other processes (Bridle & Vines 2007). The utility of our imputation is dependent on local adaptation occurring, just as in genome-environment association studies, thus both approaches are useful for identifying loci subject to selective gradients when local adaptation occurs. Our imputation can be considered a heuristic to be improved by further development.

### Genotype-by-environment interactions in genome-wide association studies

Recent advances in association models have included explicit modeling of categorical G×E (Murcray *et al.* 2009; Thomas 2010; Korte *et al.* 2012; Marigorta & Gibson 2014; Li *et al.* 2014; Kooperberg *et al.* 2016; Windle 2016), but to our knowledge there are no published genome-wide association studies accounting for genotype interactions with continuous environmental gradients (a reaction norm approach, cf. Jarquín *et al.* 2014; Tiezzi *et al.* 2017). By employing a reaction norm approach to G×E (as we did here), models can be applied to prediction at new sites, which is not possible using categorical, correlated trait approaches to G×E (Falconer 1952; Korte *et al.* 2012) where sites are treated as idiosyncratic. Some of the aforementioned categorical treatments of SNP×environment interactions were used in association studies for human disease. However, many of the environmental variables that mediate genetic risk of disease are continuous in nature, such as exposure to ultraviolet radiation and tobacco smoke. Future research on local adaptation and human disease may benefit from exchange of approaches given the shared importance across disciplines of understanding the genomic basis of G×E.

### *Case study on* Arabidopsis thaliana

Our approach identified many SNPs where allelic variation was associated with rank-changing relative fitness tradeoffs along climate gradients (*e.g*. all 214 of the SNPs with strongest interaction with winter minimum temperature association for fitness, *i.e.* 0.001 quantile), loci where selective gradients may stably maintain population differentiation (Anderson *et al.* 2011b; Ågren *et al.* 2013). Studies of local adaptation genomics often find limited evidence for loci with such fitness tradeoffs (antagonistic pleiotropy). A previous study of the common garden data used here (Fournier-Level *et al.* 2011a) found that the SNPs with the strongest association with fitness in one common garden were rarely among those with the strongest associations in another garden, which the authors interpreted as evidence for conditional neutrality. However, the fact that a locus is not among the strongest associated with fitness at an individual site does not indicate the locus is neutral at that site, it may simply be under relatively weaker selection (see Figure S7 for example illustration). By contrast with previous approaches that model phenotypes at a single site, our model was explicitly focused on detecting alleles with the strongest evidence for SNP×climate interactions favoring home alleles. Thus our explicit model of G×E is more likely to detect loci with patterns indicative of antagonistic pleiotropy compared with approaches that model fitness in a single common garden at a time, approaches that do not model G×E. Nevertheless, apparent tradeoffs at the level of individual QTL or SNPs may be driven by complementary conditionally neutral mechanisms at tightly linked loci.

Local adaptation may often involve complex traits governed by many loci. Loci exhibiting antagonistic pleiotropy and loci exhibiting G×E but no tradeoffs (variance changing or conditionally neutral) may both underlie genome-level local adaptation. Note that our study, like that of Fournier-Level *et al.* (2011a) is based on association mapping, which may suffer from identification of more false positives compared with linkage mapping approaches (Hall *et al.* 2010; Anderson *et al.* 2013; Ågren *et al.* 2013). Follow-up experimental study of phenotypic effects of variation at individual loci is required to confirm the results of association mapping (Verslues *et al.* 2014; Broekgaarden *et al.* 2015). The SNPs that exhibited the strongest evidence for SNP×climate interaction effects on fitness often fell within the coding regions of strong candidate genes based on known roles in environmental responses, suggesting our approach is a useful for identifying loci underlying local adaptation.

We found evidence that SNP×climate interaction effects on fitness were enriched in genic regions, suggesting that our model captured a signal of local adaptation rather than population structure. We found that enrichments in genic SNPs only emerged after using a mixed model to control for the putative effects of population structure (genome-wide similarity), suggesting that the genic-enriched patterns of divergence we modeled were not simply associated with overall patterns of among-population divergence. This enrichment is consistent with other findings in Arabidopsis (Hancock *et al.* 2011; Lasky *et al.* 2012b) and other species (Coop *et al.* 2009; Fumagalli *et al.* 2011; Lasky *et al.* 2015, but see Pyhäjärvi *et al.* 2013). We do not interpret this enrichment as indicating that changes in amino acid sequences are more important than regulatory evolution in local adaptation, but rather as supporting the hypothesis that local adaptation is more likely to involve sequence evolution near genes as opposed to locations farther from genes, where many intergenic SNPs are found.

We found evidence that loci we identified as candidates were enriched in evidence for partial selective sweeps (PHS and iHS statistics), suggesting that recent sweeps in particular environments are an important mode of local adaptation (Voight *et al.* 2006; Toomajian *et al.* 2006). These local sweeps may be expected based on the range dynamics of Arabidopsis, which has colonized much of its Eurasian range following the retreat of glaciers (Sharbel *et al.* 2000), a process that likely involved recent local adaptation. It is important to note that extended haplotype patterns suggestive of partial sweeps may occur at the shoulders (away from causal loci) of complete sweeps (Schrider *et al.* 2015), thus caution is warranted in attributing our observed PHS and iHS enrichment to localized sweeps versus global sweeps at nearby loci.

We found significant overlap between SNPs associated with G×E for fitness along growing season cold gradients and SNPs associated with G×E for flowering time across growing season temperature treatments (Li *et al.* 2014). This concordance suggests that variants causing flowering time plasticity drive changes in fitness across temperature gradients. Thus the evolution of temperature-responsive plasticity in flowering time may be a mechanism of local adaptation to environments that differ in growing season temperatures. For organisms inhabiting seasonal environments, timing of the life cycle can have large impacts on fitness. Previous common garden experiments have provided strong evidence that flowering time is a central trait involved in local adaptation (Hall & Willis 2006; Franks *et al.* 2007; Keller *et al.* 2012; Lowry *et al.* 2014) with molecular study further supporting the role of flowering time (Stinchcombe *et al.* 2004; Caicedo *et al.* 2004; Shindo *et al.* 2005; Lovell *et al.* 2013) and the role of plasticity (Fraser 2013; Lasky *et al.* 2014) in local adaptation to climate.

Though there was overlap with signal identified by previous approaches using the same data (Hancock *et al.* 2011; Lasky *et al.* 2012b, 2014), overlap was generally weak, indicating our approach identified distinct loci. This weak overlap is likely due to the different statistical approaches used (partial Mantel, linear mixed models, and redundancy analysis) and the additional information contained in fitness data. Furthermore, the common garden data contain signatures of processes that differ to an unknown degree from processes generating genome-environment associations. Here, the common garden data represented one generation of fitness variation among sites, whereas spatial genomic patterns accumulated due to multiple evolutionary mechanisms acting over thousands of generations.

## Conclusions

Genome-wide association studies are a promising approach for identifying the genomic basis of local adaptation and G×E. Given that many selective gradients driving local adaptation are continuous in nature, reaction norm models of G×E across multiple common gardens are a key tool for quantifying mechanisms of local adaptation. Various approaches such as genome-wide expression profiling (Des Marais *et al.* 2013), metabolomics (Meijón *et al.* 2016), and ecophysiology (Keller *et al*. 2011) are useful for uncovering integrated mechanisms of local adaptation Future approaches that use a principled basis for quantitative synthesis of patterns in multiple data types (e.g. Levy Karin *et al.* 2017) may enhance our ability to characterize mechanisms of adaptation across levels of biological organization.

## Acknowledgements

We thank David Lowry, Thomas Juenger, and six anonymous reviewers for comments on earlier versions of this manuscript.

## Data Accessibility

All data were previously published, including fitness (Data Dryad package Fournier-Level *et al.* 2011b), environment (Data Dryad package Lasky *et al.* 2012a), and SNPs (Horton *et al.* 2012, available at http://bergelson.uchicago.edu/wp-content/uploads/2015/04/call_method_75.tar.gz). Simulation data previously published in Forester *et al.* (2016) are archived in Data Dryad package (Forester *et al.* 2015). New simulation data, and code for simulations and GWAS are included in an online supplemental file with this manuscript.

## Author contributions

JRL designed research and led paper writing. All authors conducted analyses and contributed to writing.

## Supporting information

### Simulations

Forester *et al.* (2016) previously simulated local adaptation in a square two-dimensional 1024 x 1024 grid-cell landscape along a continuous environmental gradient, using the program CDPOP v1.2 (Landguth & Cushman 2010). Forester *et al.* (2016) simulated 5,000 diploid individuals with 100 bi-allelic loci, with one locus under selection and 99 loci evolving neutrally. All loci had a 0.0005 mutation rate per generation, free recombination, and no physical linkage. The authors ran 10 Monte Carlo replicates of the simulation for 1,250 generations, using the first 250 generations as a burn-in, with no selection imposed, to establish a spatial genetic pattern.

Selection varied linearly along an environmental gradient, such that *AA* and *aa* genotypes were favored at different ends of the gradient (North and South, Fig. S1). The selection strength of *s*=0.10 at extreme ends of the gradient was mediated through density-independent mortality determined by an individual’s genotype at the selected locus, where *AA* experienced 0% mortality at the North extreme and *s* mortality at the South extreme, while *aa* genotypes experienced the opposite selection gradient. *Aa* genotypes experienced uniform selection of *s*/2 across the gradient. The number of offspring produced from mating (fitness) was determined from a Poisson distribution (λ = 4), These same demographic parameters were used for common garden simulations conducted for the present paper.

In Forester *et al.* (2016), this level of fecundity produced an excess of individuals each generation. Carrying capacity of the landscape surface was 5,000 individuals, and excess new individuals were discarded once all 5,000 locations became occupied.

Mating pairs of hermaphroditic individuals and dispersal locations of offspring were chosen using a random draw from the inverse-square probability function of distance, truncated at a distance equal to *d,* the proportion of a landscape edge. We tested three values of *d*: 0.03, 0.1, and 0.25 (*i.e.* truncated at 31, 102, and 256 pixels, respectively). Individuals near landscape edges were unable to disperse or mate with individuals beyond the edge such that boundaries were not periodic.

**Tables S1-S4 (attached csv files)**. List of Arabidopsis genes within 1 kb of SNPs in the lower 0.001 quantile for p-values for SNP×environment interactions for each climate variable, including imputed observations and accounting for kinship.

**Figure S1.**
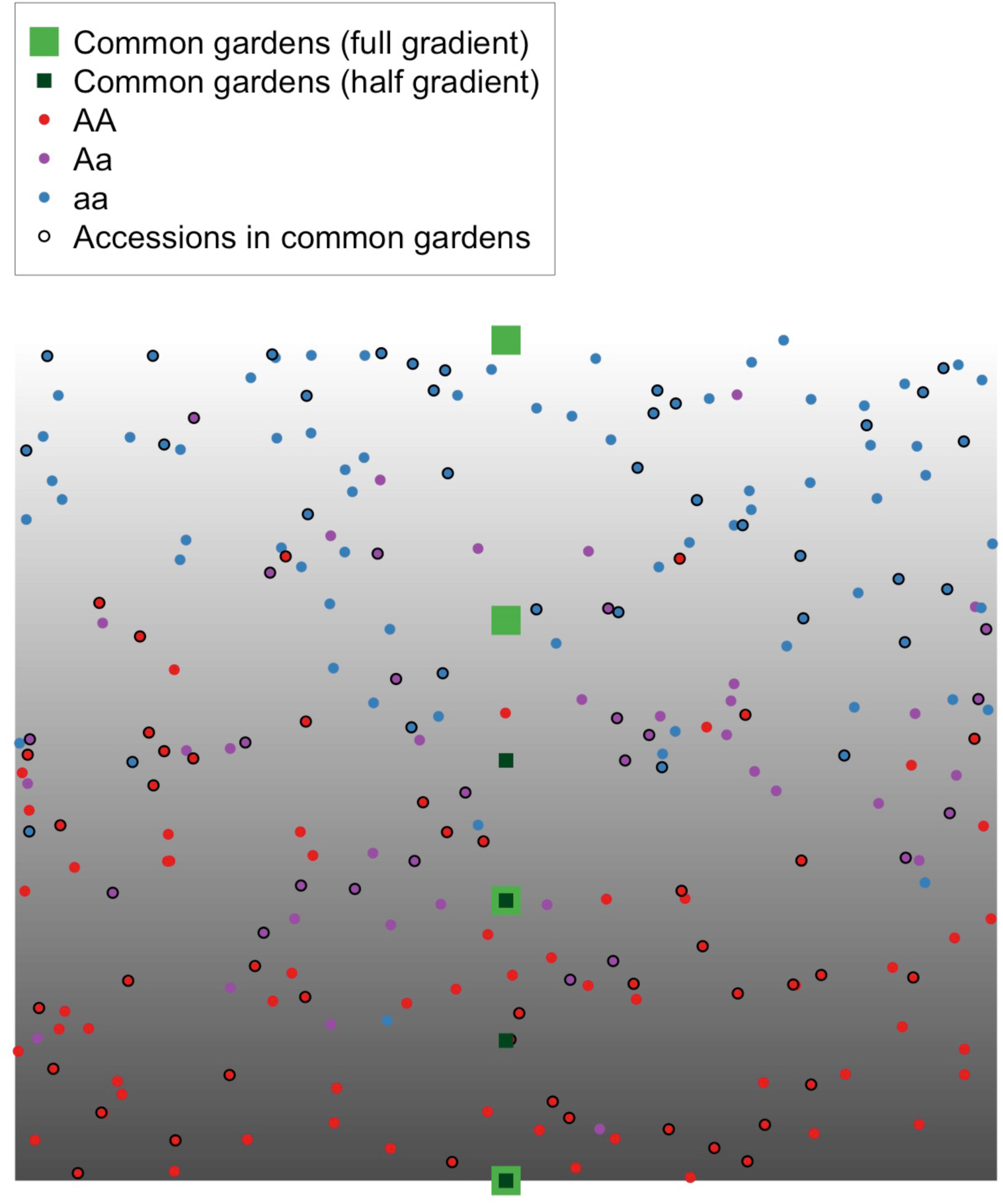
Example outcome of selection on a simulated landscape under moderate dispersal (maximum dispersal of 10% of the landscape surface per generation), with selective gradient (grayscale background), 250 randomly sampled accessions (small circles), and four common gardens (green squares). 100 accessions were simulated in the four common gardens (i.e. reciprocal transplant) covering the full gradient (large light green squares) and in four gardens covering only half the gradient (smaller dark green squares).

**Figure S2.**
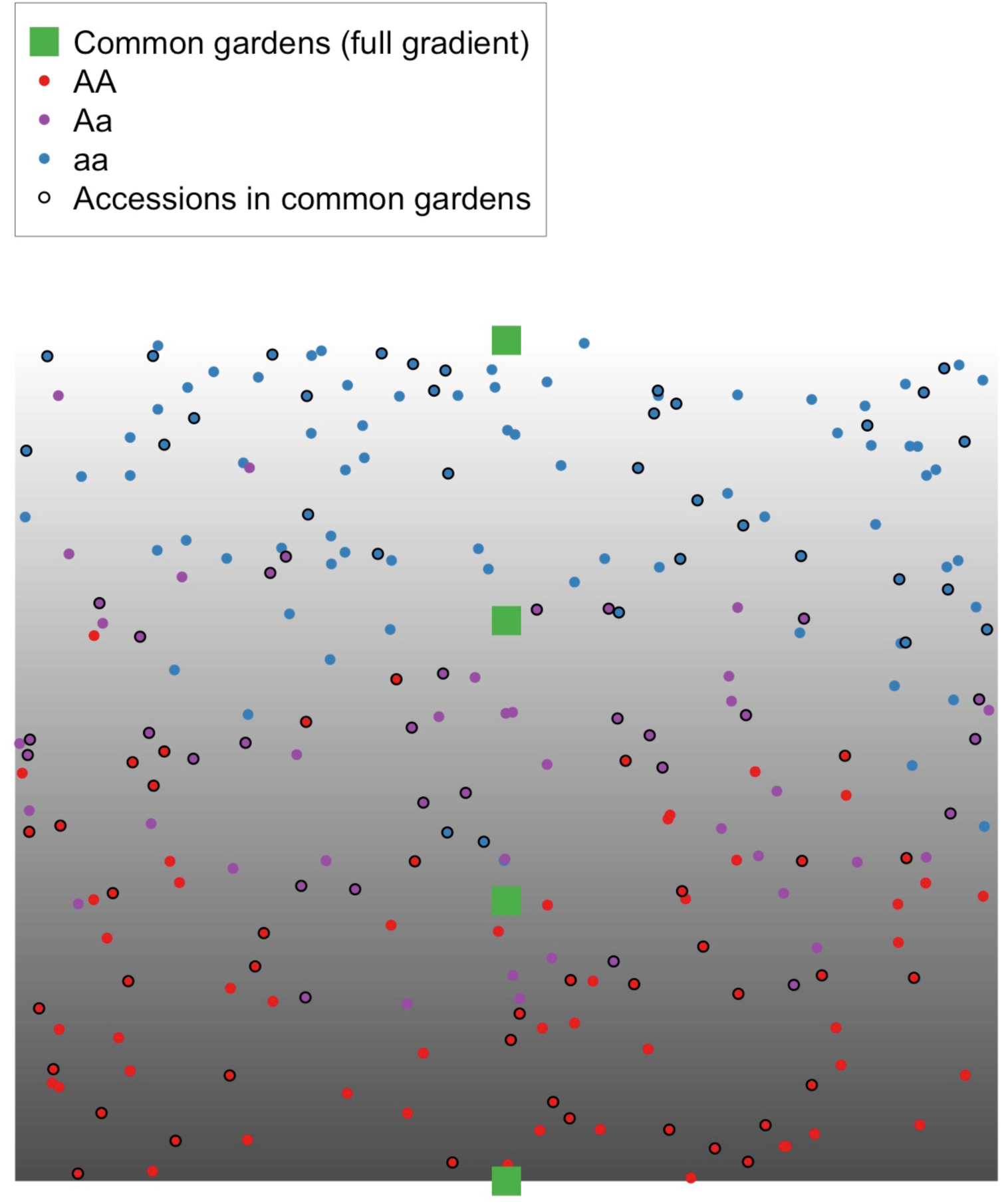
Example outcome of selection on a simulated landscape under low dispersal (maximum dispersal of 3% of the landscape surface per generation), with selective gradient (grayscale background), 250 randomly sampled accessions (small circles), and four common gardens (green squares). 100 accessions were simulated in the four common gardens (i.e. reciprocal transplant) covering the full gradient (large light green squares).

**Figure S3.**
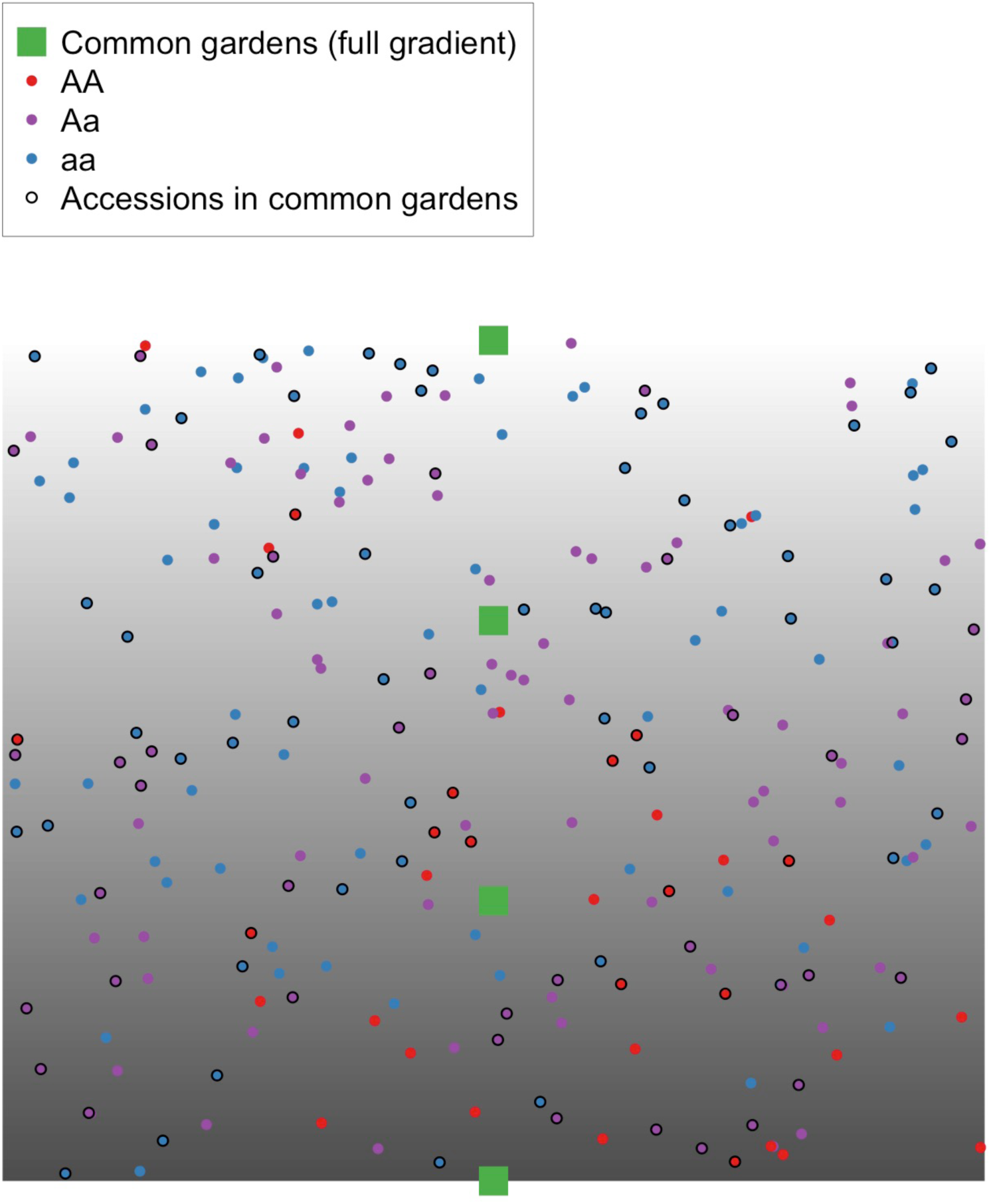
Example outcome of selection on a simulated landscape under high dispersal (maximum of 25% of the landscape surface per generation), with selective gradient (grayscale background), 250 randomly sampled accessions (small circles), and four common gardens (green squares). 100 accessions were simulated in the four common gardens (i.e. reciprocal transplant) covering the full gradient (large light green squares).

**Figure S4.**
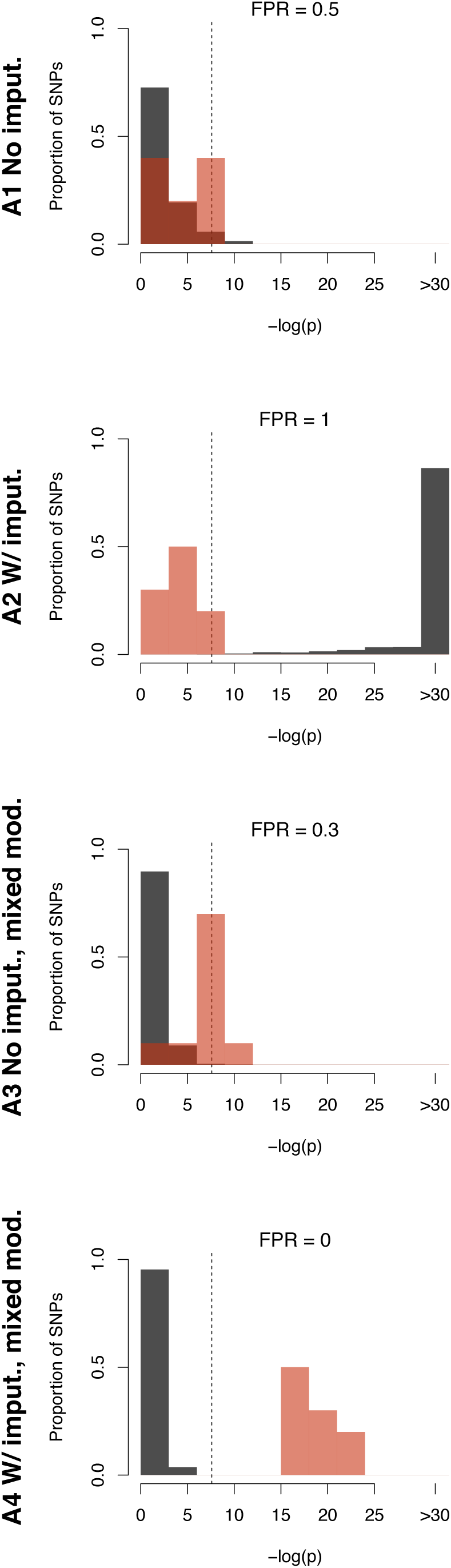
Comparison of inferred causal (red) and neutral (black) SNP associations with G×E for fitness across 10 replicate simulations for moderate dispersal with common gardens covering only half the gradient. Approaches used (row 1) no imputation and no random effects, (row 2) imputation but no random effects, (row 3) mixed models that used only observations from four common gardens, or (row 4) mixed models combining imputed observations of relative fitness in home environments with common garden observations. False positive rate (FPR) is indicated, calculated as the proportion of simulations where a neutral SNP had the lowest p-value. Each simulation had 1 causal SNP and 99 neutral SNPs; plots show aggregate distributions for all SNP by simulation combinations (*i.e.* total of 10 causal and 990 neutral SNPs). Y-axes show the proportion of SNPs in each category (causal or neutral) falling into a given p-value bin. For reference, dashed line indicates a strict Bonferroni cutoff for alpha = 0.05, –log(0.05/100) = 7.6.

**Figure S5.**
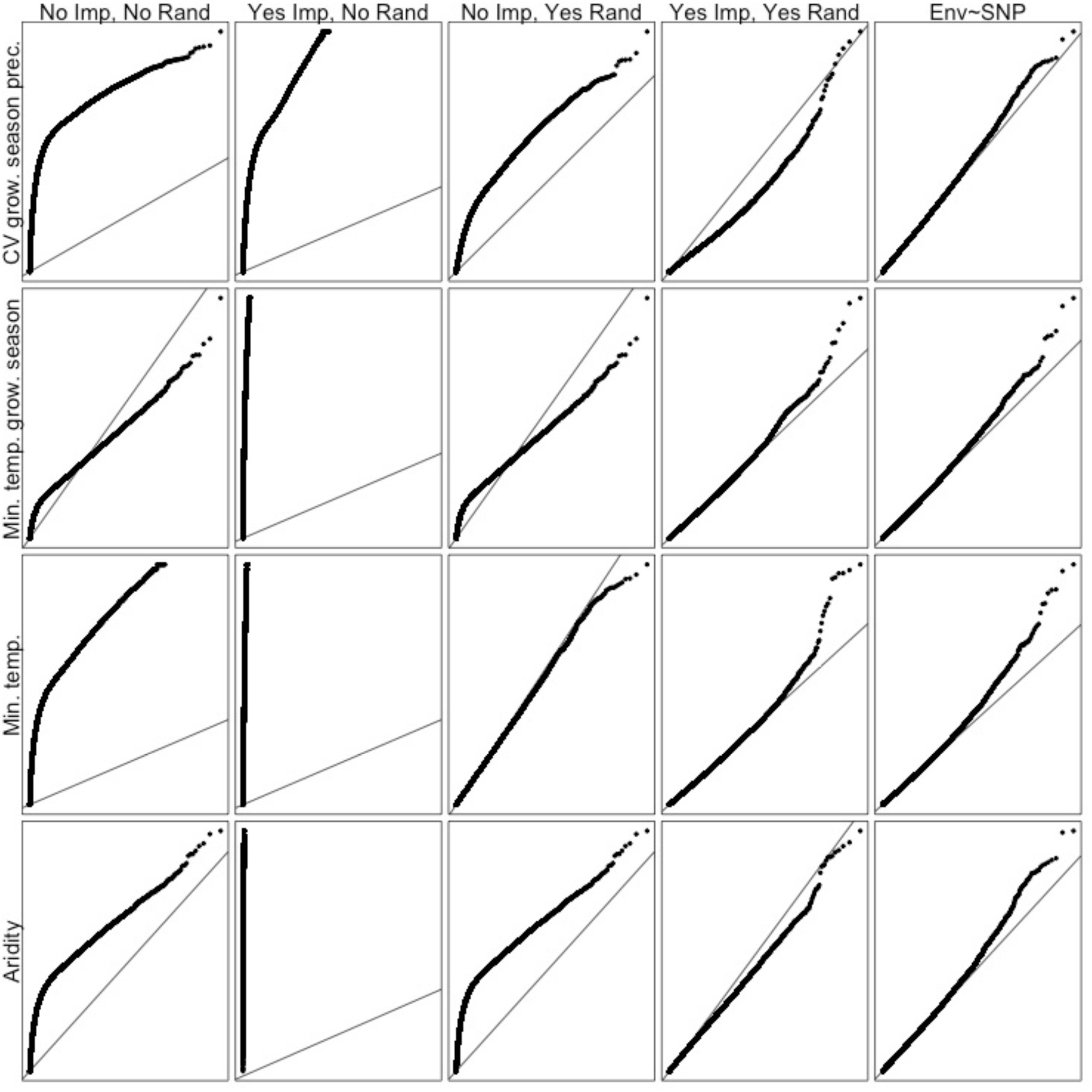
Quantile-quantile plots of p-value distributions for four approaches to calculating genome-wide SNP×environment associations with fitness (columns 1-4) and one genome-environment approach (column 5, results from Lasky *et al*. 2014), using published data on *Arabidopsis thaliana*. X-axes show expected –log_10_(p) and y-axes show observed –log_10_(p). Approaches used (column 1) no imputation and no random effects, (column 2) imputation but no random effects, (column 3) mixed models that used only observations from four common gardens, or (column 4) mixed models combining imputed observations of relative fitness in home environments with common garden observations. In the fifth column, environment is tested as a function of SNP alleles in a typical mixed model GWAS approach where environment is substituted for phenotype (Lasky *et al.* 2014).

**Figure S6.**
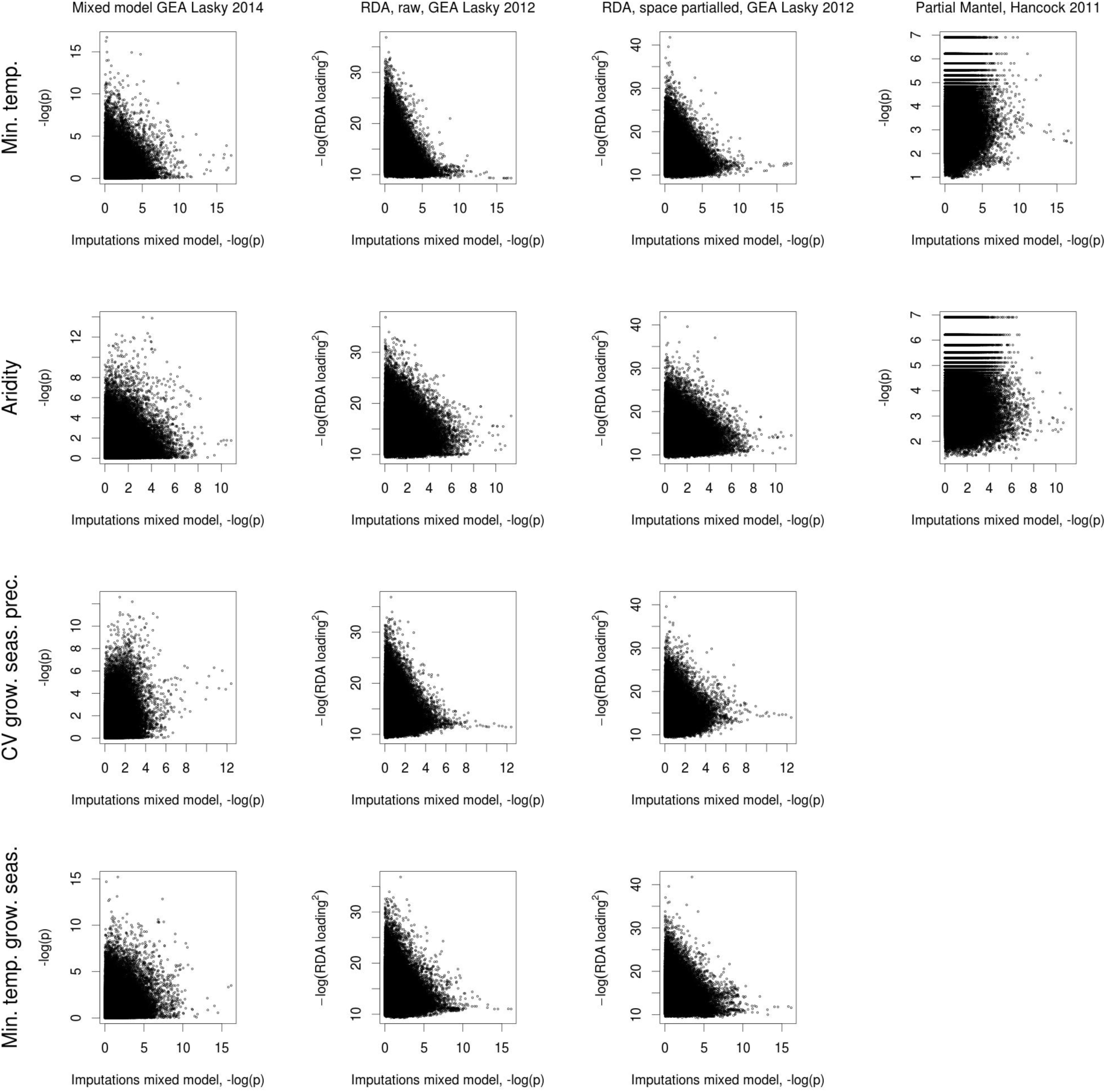
Comparison of published genome-environment association approaches to identifying loci underlying local adaptation in *Arabidopsis thaliana* (Hancock *et al.* 2011, Lasky *et al.* 2012, Lasky *et al.* 2014). Each *x*-axis shows results from the imputation approach developed in this paper, using a mixed model accounting for kinship.

**Figure S7.**
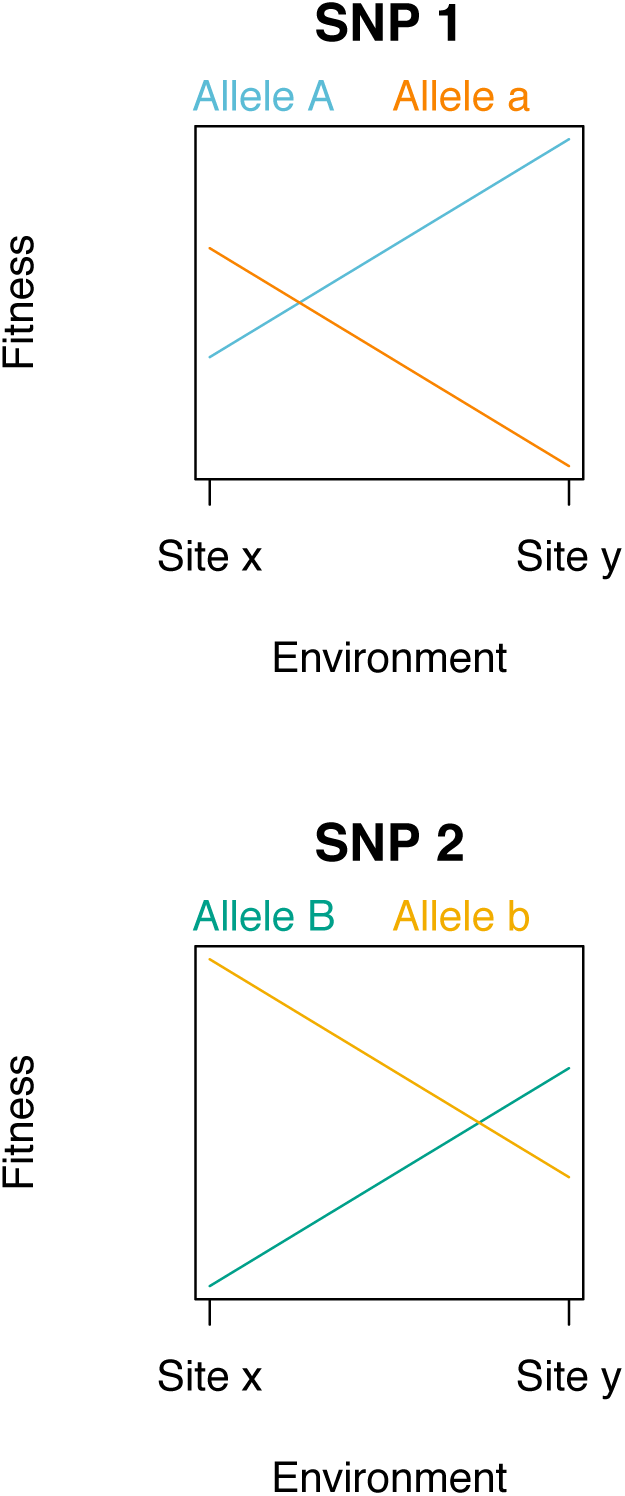
Example of fitted effects on fitness of allelic variation at two SNPs (arbitrary labeled 1 and 2) across two common gardens (arbitrary labeled x and y). In this scenario, both SNPs exhibit antagonistic pleiotropy (rank changing fitness tradeoffs) across the environmental gradient. However, each SNP has relatively weaker effects on fitness in one environment. A test of whether an individual SNP shows the strongest associations with fitness in an individual garden (*e.g.* SNP 1 at site x) might incorrectly infer conditional neutrality because other markers show stronger associations (SNP 2 at site x).

**Table S5.**
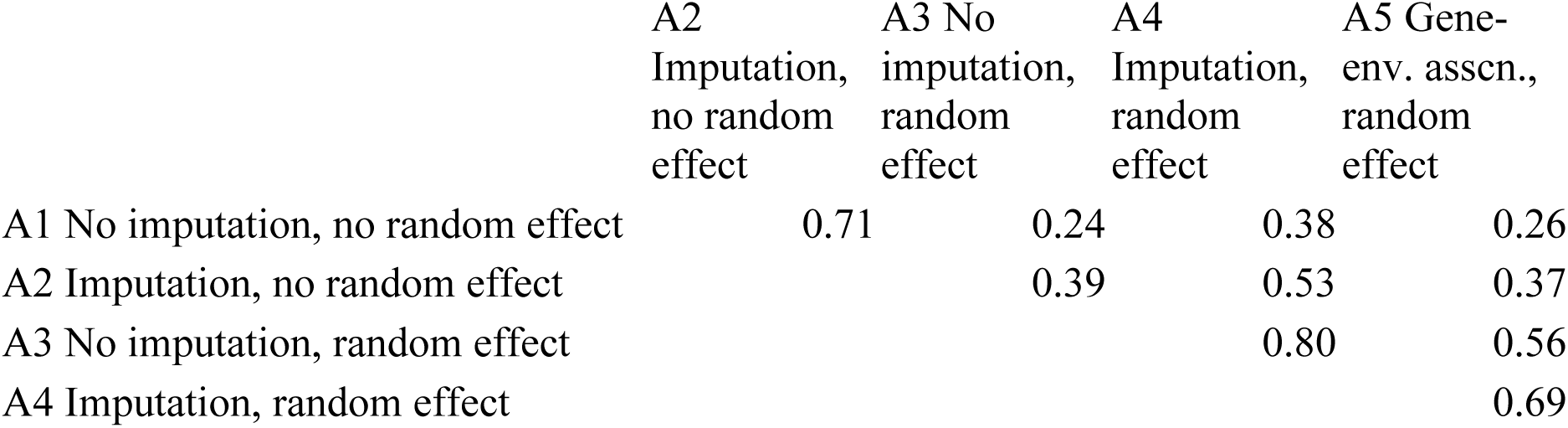
Comparison of scenarios under which five approaches studied in simulation identify true causal SNPs. Cells represent the correlation coefficient between each approach’s success at identifying the causal SNP as having the lowest p-value, across 10 different simulations across 3 dispersal levels (i.e. correlation between success for 30 total simulations).

**Table S6.** Rank correlation among SNPs for SNP×environment effect p-values comparing three tested approaches (Approaches 2-4) with Arabidopsis.

|  | *A3*<br>*Random*<br>*effect, no*<br>*imputation* | *A4*<br>*Random*<br>*effect,*<br>*imputation* |
| --- | --- | --- |
| **Min. temp. growing season** |  |  |
| *A2 No random effect, imputation* | 0.396 | -0.001 |
| *A3 Random effect, no imputation* |  | 0.041 |
| **Aridity** |  |  |
| *A2 No random effect, imputation* | 0.318 | 0.008 |
| *A3 Random effect, no imputation* |  | -0.032 |
| **Min. temp. coldest month** |  |  |
| *A2 No random effect, imputation* | -0.021 | -0.012 |
| *A3 Random effect, no imputation* |  | 0.104 |
| **CV growing season prec.** |  |  |
| *A2 No random effect, imputation* | 0.178 | 0.090 |
| *A3 Random effect, no imputation* |  | -0.006 |

**Table S7.** Overlap (one-tailed permutation tests) and rank correlations (Spearman’s rho) between our approach (Approach 4, mixed-model with imputation, climate variables give row labels at left) and published approaches to identifying SNPs associated with local adaptation in Arabidopsis. Citations are given for the original publication of previous approaches. For partial Mantel and RDA, we used absolute value of SNP scores to rank SNPs. Empty cells indicate climate variables not tested in Hancock *et al*. (2011).

| Climate var. | Mixed model, overlap in 0.01 tail strongest associations<br>Lasky et al. 2014 | Mixed model, rank correlation between associations<br>Lasky et al. 2014 | First axis of RDA, overlap in 0.01 tail strongest associations<br>Lasky et al. 2012 | First axis of RDA, rank correlation between associations<br>Lasky et al. 2012 | First axis of RDA after removing spatial structure, overlap in 0.01 tail strongest associations<br>Lasky et al. 2012 | First axis of RDA after removing spatial structure, rank correlation between associations<br>Lasky et al. 2012 | Partial Mantel overlap in 0.01 tail strongest associations<br>Hancock et al 2011 |
| --- | --- | --- | --- | --- | --- | --- | --- |
| Min. temp. coldest month | <0.0001 | 0.0416 | <0.0001 | -0.2126 | <0.0001 | -0.0575 | 0.015 |
| Aridity | <0.0001 | 0.0573 | <0.0001 | -0.0153 | 0.1137 | 0.0127 | <0.000 |
| CV growing season prec. | <0.0001 | 0.1871 | 0.9623 | -0.0282 | 0.9959 | 0.0303 |  |
| Min. temp. growing season | <0.0001 | 0.0794 | 0.0556 | -0.0928 | <0.0001 | -0.1319 |  |

